# GM1 mediates the formation and maintenance of cytotoxic Aβ oligomers

**DOI:** 10.1101/2021.07.22.453392

**Authors:** Dong Yan Zhang, Jian Wang, Rebecca M. Fleeman, Madison K. Kuhn, Matthew T. Swulius, Elizabeth A. Proctor, Nikolay V. Dokholyan

## Abstract

The aggregation of amyloid beta (Aβ) peptide is associated with Alzheimer’s disease (AD) pathogenesis. Cell membrane composition, especially monosialotetrahexosylganglioside (GM1), is known to promote the formation of Aβ fibrils, yet little is known about the roles of GM1 in the early steps of Aβ oligomer formation. Here, by using GM1-contained liposomes as a mimic of neuronal cell membrane, we demonstrate that GM1 is a critical trigger of Aβ oligomerization and aggregation. We find that GM1 not only promotes the formation of Aβ fibrils, but also facilitates the maintenance of Aβ oligomers on liposome membranes. We structurally characterize the Aβ oligomers formed on the membrane and find that GM1 captures Aβ by binding to its arginine-5 residue. To interrogate the mechanism of Aβ oligomer toxicity, we design a new liposome-based Ca^2+^-encapsulation assay and provide new evidence for the Aβ ion channel hypothesis. Finally, we conduct cell viability assay to determine the toxicity of Aβ oligomers formed on membranes. Overall, by uncovering the roles of GM1 in mediating early Aβ oligomer formation and maintenance, our work provides a novel direction for pharmaceutical research for AD.

## INTRODUCTION

Alzheimer’s disease is the most common neurodegenerative disorder, responsible for 60–70% of dementia cases (Burns and Iliffe, 2009). The economic burden associated with caring for nearly 50 million people worldwide with AD is estimated in the hundreds of billions of dollars annually (Vos et al., 2016). The pathophysiology of AD is characterized by the accumulation of extracellular plaques, predominantly composed of amyloid beta (Aβ), and cytoplasmic neurofibrillary tangles, mostly composed of tau protein. Despite multiple proposed hypotheses (Alves et al., 2015; Bartzokis, 2011; Cataldo et al., 2010; Deane and Zlokovic, 2007; Francis et al., 1999; Hardy and Allsop, 1991; Kandimalla et al., 2016; Miklossy, 2011; Mudher and Lovestone, 2002; Pisa et al., 2015; Su et al., 2008), the molecular etiology of the disease remains a mystery. One of the oldest and most central hypotheses, the *amyloid cascade hypothesis*, posits that the accumulation of Aβ proteins in the brain is the primary cause of AD pathogenesis, leading to tau pathology, neuroinflammation, synapse loss, and ultimately neuron death (Hardy, 2006; Hardy and Higgins, 1992; Iversen et al., 1995; Jakob-Roetne et al., 2009; Tanzi and Bertram, 2005).

The *amyloid cascade hypothesis* was subsequently revised to *A*β *oligomers cascade hypothesis* (Bernstein et al., 2009; Butterfield and Lashuel, 2010; Glabe, 2008; Matsumura et al., 2011; Quist et al., 2005), stating that small Aβ oligomers (DeToma et al., 2012; Haass and Selkoe, 2007) rather than fibrils (Klein et al., 2001; Lambert et al., 2001) are the main toxic species, is an alternative to the original amyloid cascade hypothesis. The oligomer hypothesis has been gaining significant momentum, since numerous studies have shown that Aβ oligomers are toxic to primary neurons, inhibit hippocampal long-term potentiation, and cause memory impairment in rat or mouse models (Karran et al., 2011; Martinez Hernandez et al., 2018). Mounting evidence, stemming from other neurodegenerative diseases, also support the oligomer hypothesis. For example, in amyotrophic lateral sclerosis, soluble superoxide dismutase (SOD1) oligomers (Choi and Dokholyan, 2021; Proctor et al., 2015; Redler et al., 2014) of disease-associated proteins, rather than insoluble aggregates (Zhu et al., 2018), are shown to be responsible for cytotoxicity. Insoluble SOD1 aggregates have been found to be protective against neuronal toxicity, potentially due to competition with soluble oligomers (Bieschke et al., 2011; Zhu et al., 2018).

The mechanism of Aβ oligomer toxicity is unknown, but several studies (Anekonda et al., 2011; Bhowmik et al., 2015; Jang et al., 2007, 2010; Quist et al., 2005) show that Aβ oligomers may disrupt the plasma membrane to upset ionic (especially calcium) homeostasis. A number of mechanisms by which Aβ oligomers induce cell membrane disruption have been proposed, including membrane thinning (Dante et al., 2008), excessive membrane tabulation (Pandey et al., 2011; Varkey et al., 2010), membrane extraction through amyloid–lipid co-aggregation (Hellstrand et al., 2013; Reynolds et al., 2011), and formation of ion channels to disrupt Ca^2+^ homeostasis (Arispe et al., 1993a; Shirwany et al., 2007). Membrane components can also promote the conformational change of Aβ from α-helix-rich structures to β-sheet-rich structures (Ikeda et al., 2011) and play a determining role in the uptake of Aβ oligomers (Di Scala et al., 2016). Since cellular membranes are highly heterogeneous, containing many constituents, deciphering which cellular membrane components are catalyzing Aβ cytotoxic action is challenging to untangle. Specifically, AD patients have much higher amounts of monosialotetrahexosylganglioside (GM1), a glycosphingolipid found in neuronal cell membranes (Yagi-Utsumi et al., 2010), than cognitively normal patients in the cerebrospinal fluid (Okada et al., 2007). GM1 clusters (Fernández-Pérez et al., 2017; Ikeda et al., 2011; Tachi et al., 2019; Yagi-Utsumi et al., 2010) are found to facilitate the α-to-β conformational transition of Aβ40. GM1-bound Aβ has been found in early pathological changes of the AD brain (Kakio et al., 2002). While there is strong evidence that Aβ binds to GM1 and acts as a seed to further promote the formation of Aβ fibrils (Yanagisawa et al., 1995), the influence of GM1 on the formation of soluble Aβ oligomers, the most toxic form, and whether GM1 is involved in the pathways responsible for Aβ oligomer toxicity, remains a mystery.

Here, we employ *in vitro, in silico*, and cellular studies to interrogate the interaction between GM1 and Aβ oligomers, and to uncover the mechanisms driving the assembly and toxicity of Aβ oligomers (Figure 1). We utilize liposomes, artificial vesicles (0.05 µm to 5 µm in diameter (Mufamadi et al., 2011)) formed from phospholipid bilamellar membranes, to determine the interactions between Aβ and lipid membranes in the presence of GM1. We find that GM1 promotes the maintenance of Aβ oligomers. Using molecular dynamics simulations, we observe the interaction between Aβ and GM1 and identify the most critical residue, R5, involved in the binding between Aβ and GM1. We develop a fluorescence assay to interrogate the mechanism of Aβ disrupting the membrane and also uncover the role of various membrane constituents on Aβ aggregation. This fluorescence assay provides new evidences supporting the formation of Aβ ion channels. We use mass spectrometry to characterize purified Aβ oligomers formed on liposomes, and find that these membrane-associated Aβ oligomers are composed of 16-21 monomers, rich in β-sheet. Finally, we perform cellular studies to demonstrate the neuronal toxicity of membrane-associated Aβ oligomers. Overall, our results provide new insights into the mechanisms of Aβ aggregation by revealing the role of GM1 in mediating early Aβ oligomer formation and maintenance. Uncovering the mechanisms of Aβ aggregation and determining critical cellular players responsible for downstream neurotoxicity is essential for the development of novel pharmaceutical strategies to treat AD.

**Figure 1.**
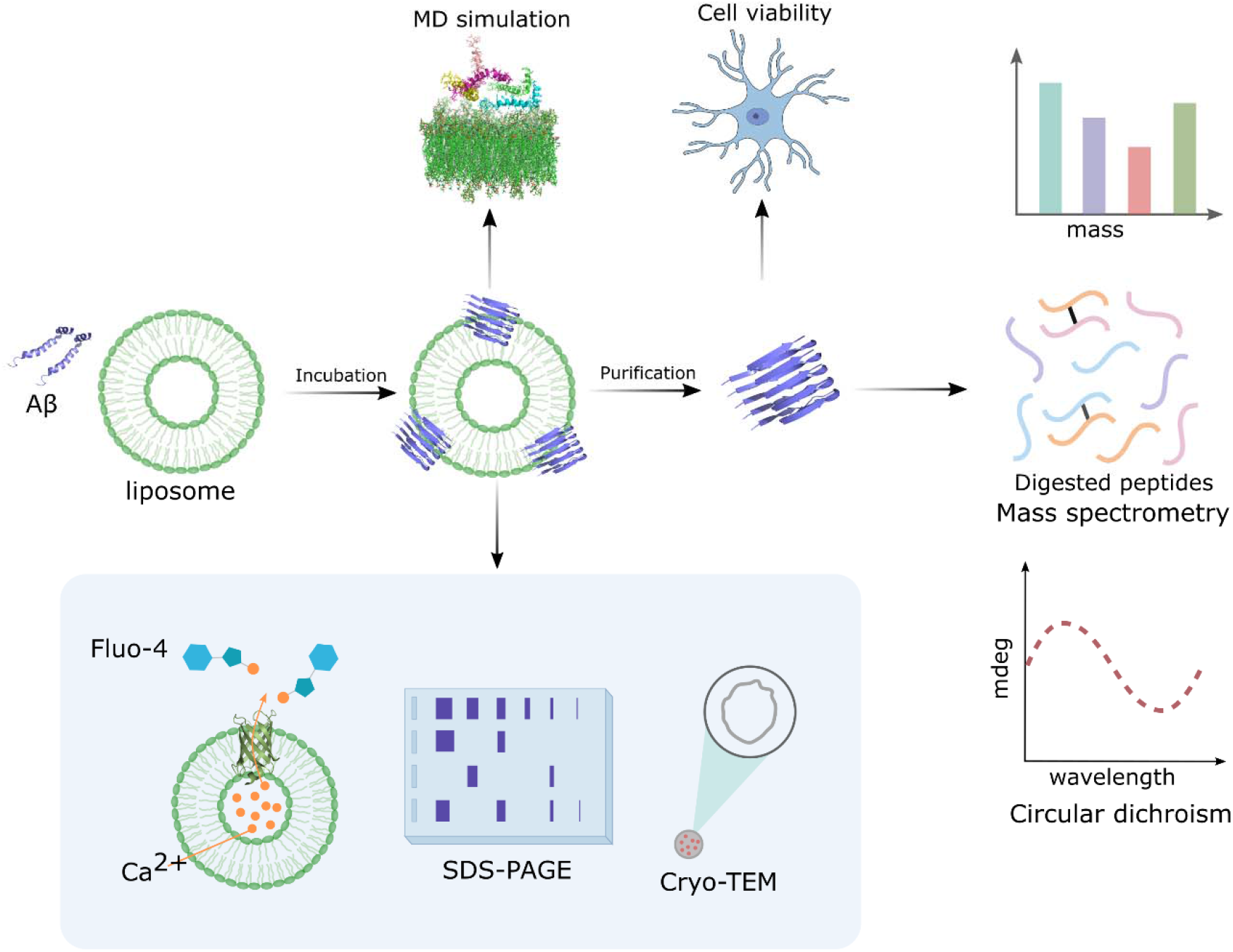
Schematic of the hypothesis and the design. We incubate Aβ with GM1 liposomes to determine the effect of GM1 membrane on the formation of Aβ fibrils and oligomers. We perform MD simulation to explore the interaction between Aβ and GM1 membrane. We purify the Aβ oligomers formed on liposomes membrane, determine the toxicity, and characterize the structure with CD and mass spectrometry. We design a calcium-encapsulation assay to provide new evidences for the Aβ ion channel hypothesis. We also determine the effect of Aβ on the morphology of liposomes membrane using Cryo-TEM.

## RESULTS

### GM1 liposomes promote Aβ fibril formation and facilitate Aβ oligomer maintenance

GM1 clusters on liposome membrane promote the formation of Aβ fibrils (Kakio et al., 2001). Do GM1 clusters promote the formation of Aβ oligomers? To answer this question, we incubate Aβ peptides in the presence and absence of liposomes consisting of GM1, sphingomyelin, and cholesterol (GM1 liposomes) for 0.5 h, 4 h, 24 h, and 72 h, respectively and then perform 10% SDS-PAGE and silver staining (Figure 2A-2D). We use cholesterol and sphingomyelin in liposomes because they are the indispensable components of GM1 clusters. We find that in the absence of GM1 liposomes, Aβ oligomers of different molecular weights are formed within 0.5 h, and the amount of Aβ oligomers species then reduces with the increase of time. After 72 h, there are nearly no Aβ oligomers. We speculate that the disappeared oligomers have formed larger assemblies, such as amyloid fibrils (Figure 2E). In the presence of GM1 liposomes, we still observe abundant Aβ oligomers species formed within 0.5 h, and the amount of oligomer species reduces even more rapidly than when the Aβ peptides are incubated without GM1 liposomes. However, after 72 h, we still observe a few clear Aβ oligomer bands (30, 35, 40, 45, 60 kDa), indicating that some oligomers are arrested in their state, thus suggesting some distinct oligomer species from those formed without GM1. Thus, GM1 can promote the formation of fibrils by catalyzing oligomers to form fibrils, but GM1 can also facilitate the maintenance of certain oligomer species.

**Figure 2.**
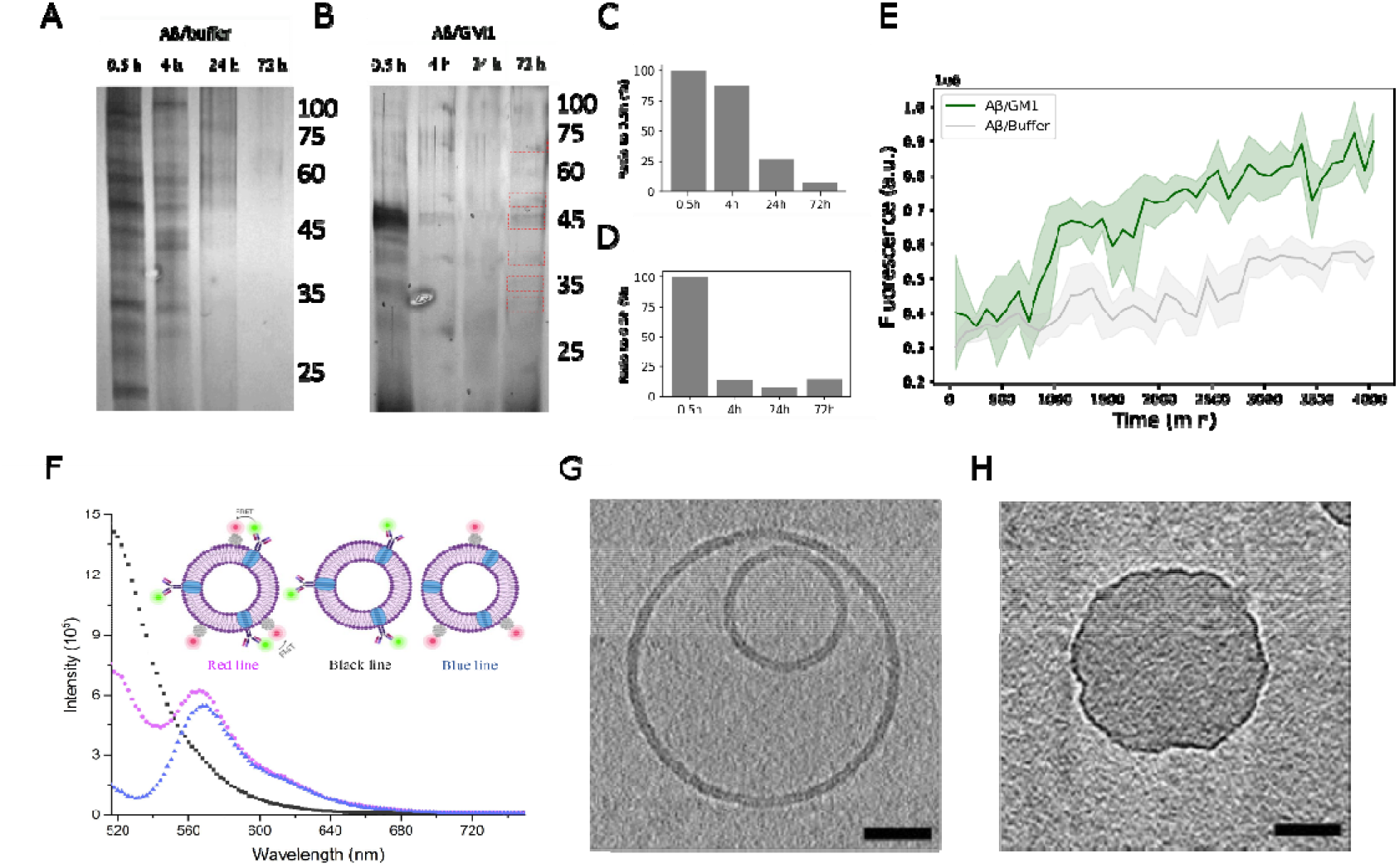
Interaction between Aβ and GM1 liposome. (A), (B) SDS-PAGE and silver staining results of Aβ oligomers formed in the presence and absence of GM1 liposomes at different time points. (C), (D) Quantitative analysis of Aβ oligomers formed in the presence and absence of GM1 liposomes at different time points. (E) The formation of Aβ fibrils is identified by ThT. (F) FRET results of GM1 and Aβ. The donor is Alexa Fluor® 488 6E10, the antibody of Aβ. The acceptor is Alexa Fluor™ 555 CTSB, a molecule that specifically binds to GM1. Red curve: Aβ, GM1 liposomes, donor and acceptor. Black curve: Aβ, GM1 liposomes and donor. Blue curve: Aβ, GM1 liposomes and acceptor. (G&H) An approximately 10-nm thick digital slices through tomograms of liposomes incubated with (G) or without Aβ (H). The lipid bilayer is visible in the untreated sample and only a monolayer is visible after treatment. Scale bars represent 50 microns.

To interrogate if the disappeared oligomers form fibrils, we determine the formation of fibrils (Figure 2E) through thioflavin T (ThT)-binding assay; ThT is a benzthiazole dye to detect Aβ fibrils (Gade Malmos et al., 2017). We incubate 80 μM Aβ with ThT in the presence and absence of GM1 liposomes and find that ThT fluorescence intensities increase with time (Figure 2E), indicating that 80 μM Aβ monomers incubated in the presence and absence of GM1 liposomes both aggregate into fibrils. Consistent with previous reports (Yanagisawa, 2005), we find that Aβ aggregates faster and forms a greater number of aggregates in the presence of GM1 liposomes, suggesting that GM1 liposomes promote the formation of Aβ fibrils. Overall, previous studies have reported that Aβ binding to GM1 is a seeding step to form larger aggregates (Yanagisawa et al., 1995), and our work further shows that (1) Aβ oligomers form rapidly in 0.5 h but then partially disappear because they begin to assemble into higher molecular weight aggregates, (2) Aβ fibrils form more rapidly in the presence of GM1, and (3) GM1 maintain some Aβ oligomer species, probably because these species may have direct interaction with GM1. These oligomers are likely to be structurally distinct from oligomers formed on-pathway to fibrils.

To confirm the direct interaction between Aβ and GM1, we utilize Förster resonance energy transfer (FRET) to detect the interaction between Alexa Fluor™ 555 CTSB labeled GM1 and Alexa Fluor® 488 6E10 labeled Aβ (Figure 2F). We add Alexa Fluor® 488 6E10, Alexa Fluor™ 555 CTSB, and a mixture of Alexa Fluor® 488 6E10 and Alexa Fluor™ 555 CTSB to the samples of Aβ incubated with GM1 liposome, respectively, to compare the fluorescence emission spectra. The fluorescence intensity peak of the donor at 518 nm reduces to approximately 51% of that of the mixture of Alexa Fluor® 488 6E10 and Alexa Fluor™ 555 CTSB, and combines with an increasing approximately 15% fluorescence intensity peak of the receptor at 566 nm. The changes of the fluorescence intensity peaks are due to the non-radiative energy transfer from Alexa Fluor® 488 to Alexa Fluor™ 555, suggesting an efficient FRET from Aβ to GM1 liposomes membrane. Thus, we speculate that the maintenance of certain Aβ oligomers species may be the result of the direct physical interaction to GM1 clusters.

In turn, Aβ oligomers also impact the morphology of GM1 membranes. We use cryo-electron tomography (cryo-TEM) to analyze the morphology of liposomes incubated with or without Aβ. In the absence of Aβ, GM1 liposomes form bilamellar vesicles with smooth surfaces (Figure 2G&H), while in the presence of Aβ, GM1 liposomes become unilamellar vesicles and the membranes of these liposomes are disordered and deformed. Thus, the presence of Aβ oligomers significantly affects the surface morphology of GM1 liposome membranes by deforming the membranes. We posit that for neurons, Aβ may interact with GM1 clusters in cell membranes (Bhowmik et al., 2015; Jang et al., 2007, 2010; Quist et al., 2005) in a similar fashion in the brain, thereby resulting in deformed membrane and, ultimately, in neurotoxicity and synaptic loss (Du et al., 2019).

### R5 of Aβ42 plays an important role in the interaction of Aβ and GM1 membranes

We perform 100 ns molecular dynamics (MD) simulations of GM1 membrane and 5 Aβ42 monomers (Figure 3A). We calculate the minimum distance between atoms of each residue of Aβ and all atoms of GM1 on the membrane. We find that the fifth residue (arginine, R5) of Aβ maintains a distance of 1-2 Å from GM1 (Figure 3C&D), suggesting that R5 stably binds GM1. Thus, we posit that R5 plays a critical role in the interaction between Aβ and GM1, specifically, the guanidine group in R5 interacting with the O11 and O12 atoms in GM1 (Figure 3B). We further computationally substitute R5 to glycine, and upon performing MD simulations we find that the average distance between G5 and GM1 increases to >10 Å (Figure 3E) and is within binding distance (<2 Å) for as little as 2% of the time (Figure 3F), indicating that the mutation of R5G disrupts the tight binding interaction between the fifth residue and GM1. This result strongly suggests the critical role of R5 in GM1-Aβ interaction.

**Figure 3.**
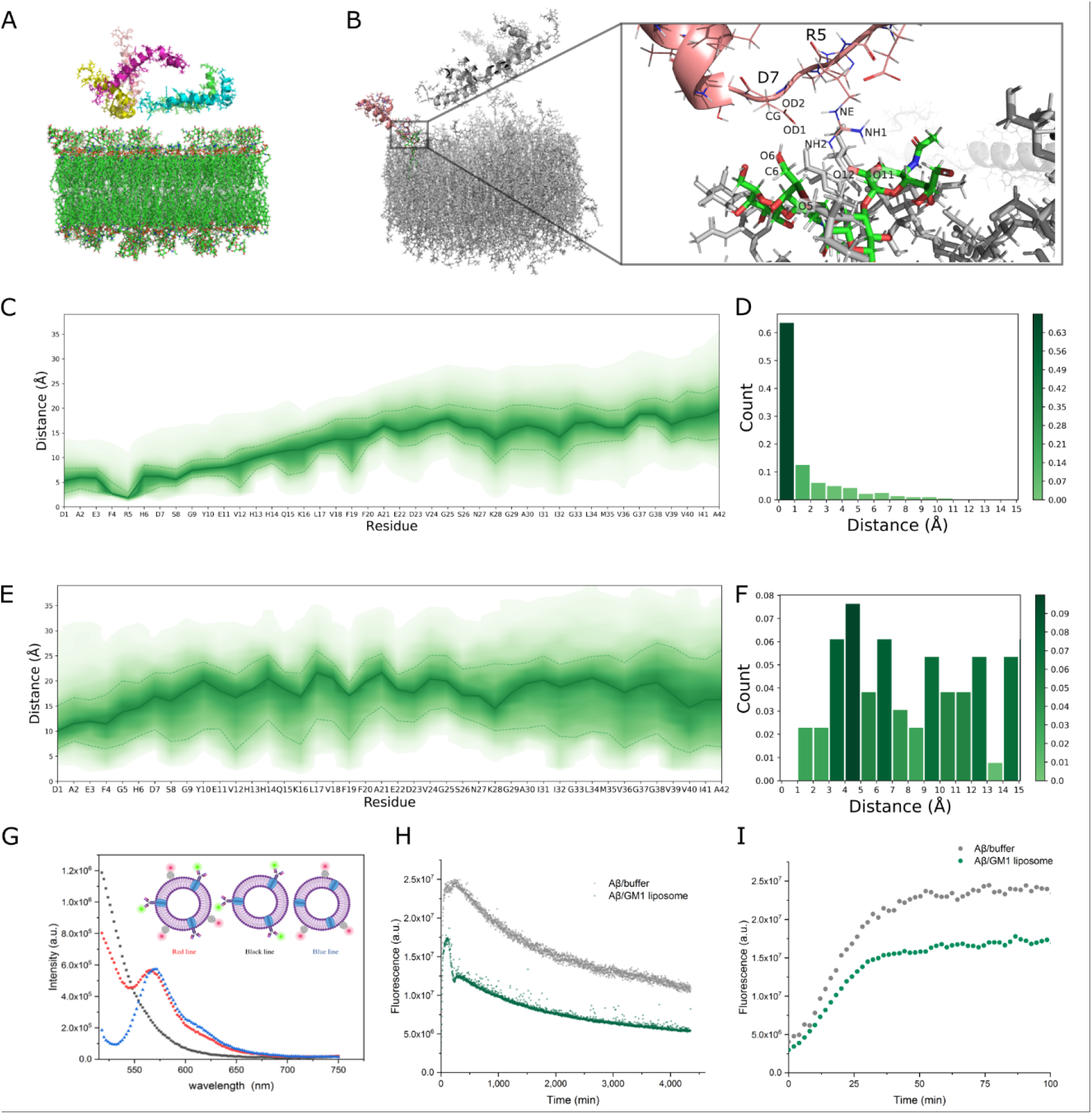
Interaction between wild type, mutant Aβ, and GM1 liposome. (A) The initial structure for the molecular dynamics simulation of GM1 membrane and 5 Aβ monomers. (B) The 3D view of the atoms in the interacting interface between R5 and GM1. (C) The distances between atoms of each residue in wild type Aβ and atoms of all GM1 in the membrane. (D) The histogram of the minimum distances between R5 in Aβ and GM1 in the membrane of all steps in the MD simulation. (E) The distances between the Aβ mutant R5G and GM1. (F) The histogram of the minimum distances between G5 and GM1. G5 is the fifth residue in the Aβ mutant R5G. (G) FRET between R5G and GM1. The donor is Alexa Fluor® 488 6E10, the antibody of Aβ. The acceptor is Alexa Fluor™ 555 CTSB, a molecule that specifically binds to GM1. Red curve: mutant Aβ, GM1 liposomes, donor and acceptor. Black curve: mutant Aβ, GM1 liposomes and donor. Blue curve: mutant Aβ, GM1 liposomes and acceptor. (H) The formation of mutant Aβ (R5G) fibrils are identified by ThT. (I) The formation of mutant Aβ (R5G) fibrils in the first 100 min.

To confirm the role of R5 experimentally, we substitute arginine with glycine and then utilize FRET to detect the interaction between the mutant Aβ and GM1 membrane (Figure 3G). The fluorescence intensity peak of the acceptor at 566 nm of the mixture of Alexa Fluor® 488 6E10 and Alexa Fluor™ 555 CTSB is similar to that of the sample containing only the acceptor. In contrast to wild type Aβ (Figure 2F), there is no energy transfer from donor to acceptor for the mutant Aβ, indicating that the mutant Aβ does not bind to the GM1 liposome membrane. To determine the effect of the mutation on the formation of fibrils, we perform the ThT-based fibril detection assay for the mutants. We find that R5G aggregates into fibrils in 100 min, then the fibrils slowly dissociate (Figure 3H&I). In comparison, the wild type Aβ fibrils do not dissociate (Figure 2E). This observation suggests that R5 also plays an important role in Aβ fibril stability. Finally, we find that compared to HEPES buffer, the presence of GM1 liposomes inhibits aggregation of the mutant Aβ into fibrils, as the sample with GM1 liposomes always feature a lower ThT fluorescence intensity.

### GM1-catalyzed channel formation by Aβ oligomers

Various mechanisms (Dante et al., 2008; Hellstrand et al., 2013; Pandey et al., 2011; Reynolds et al., 2011; Varkey et al., 2010) by which Aβ oligomers disrupt cell membranes have been proposed. Our work strongly supports the ion channel hypothesis (Kagan et al., 2002) that posits that Aβ oligomers damage neuron by forming ion channels channels (Arispe et al., 1993a; Shirwany et al., 2007). Although the hypothesis provides a biophysical mechanism for explaining the Aβ oligomers toxicity(Hane and Leonenko, 2014), it needs further validation. We develop a fluorescence assay to interrogate if the Aβ oligomers maintained by GM1 form ion channels to disrupt the plasma membrane, with the goal of uncovering the role of various membrane constituents (PC, SM, and GM1) in the disruption process. We encapsulate Ca^2+^ inside liposomes and dialyze away the excess Ca^2+^ from outside of the liposomes. In support of the ion channel hypothesis, we expect that the formation of Aβ ion channels would lead to the efflux of Ca^2+^, thus resulting in an increase of Ca^2+^ concentration outside the liposomes. We utilize a calcium sensitive dye (Fluo-4) to detect the Ca^2+^ concentration change outside of the liposomes. To verify whether we successfully encapsulated Ca^2+^ inside the liposomes, we add Triton X-100 to lyse the Ca^2+^-loaded liposomes (Supplementary Figure 1). We prepare two types of liposomes: (1) PC liposomes (L-α-Phosphatidylcholine+ sphingomyelin +cholesterol), and (2) GM1 liposomes (GM1+ sphingomyelin +cholesterol). We observe Ca^2+^ efflux after adding Triton X-100 (Supplementary Figure 1), confirming that Ca^2+^ is successfully encapsulated inside the liposomes.

We incubate varying concentrations of Aβ with Ca^2+^-encapsulated GM1 and PC liposomes. When Aβ concentration is higher than 10 μM, GM1 and PC liposomes have similar Ca^2+^ efflux, reflecting similar disruption to liposomes, yet when Aβ concentration is lower than 1 μM, GM1 liposomes have much more Ca^2+^ efflux than that of PC liposomes, indicating that GM1 liposomes are disrupted to a greater extent. (Figure 4A&B). Thus, at high Aβ concentration (>10 μM), Aβ induces leakage of both liposome types; at low Aβ concentrations, Aβ induces more leakage of GM1 liposomes than that of PC liposomes, indicating that GM1 is key to the process of Aβ-induced membrane disruption and may serve as a catalyst to promote disruption especially at low Aβ concentrations (<1 μM). To further demonstrate that PC and cholesterol are irrelevant to Ca^2+^ efflux, we also test three more types of PC liposomes with different composition proportion of PC, cholesterol, and sphingomyelin (20% PC, 40% cholesterol, 40% sphingomyelin; 40% PC, 20% cholesterol, 40% sphingomyelin; 50% PC, 10% cholesterol, 40% sphingomyelin;). We encapsulate Ca^2+^ inside these liposomes and incubate them with Aβ. We find no significant Ca^2+^ efflux (Supplementary Figure 2), indicating no Aβ-induced membrane disruption.

**Figure 4.**
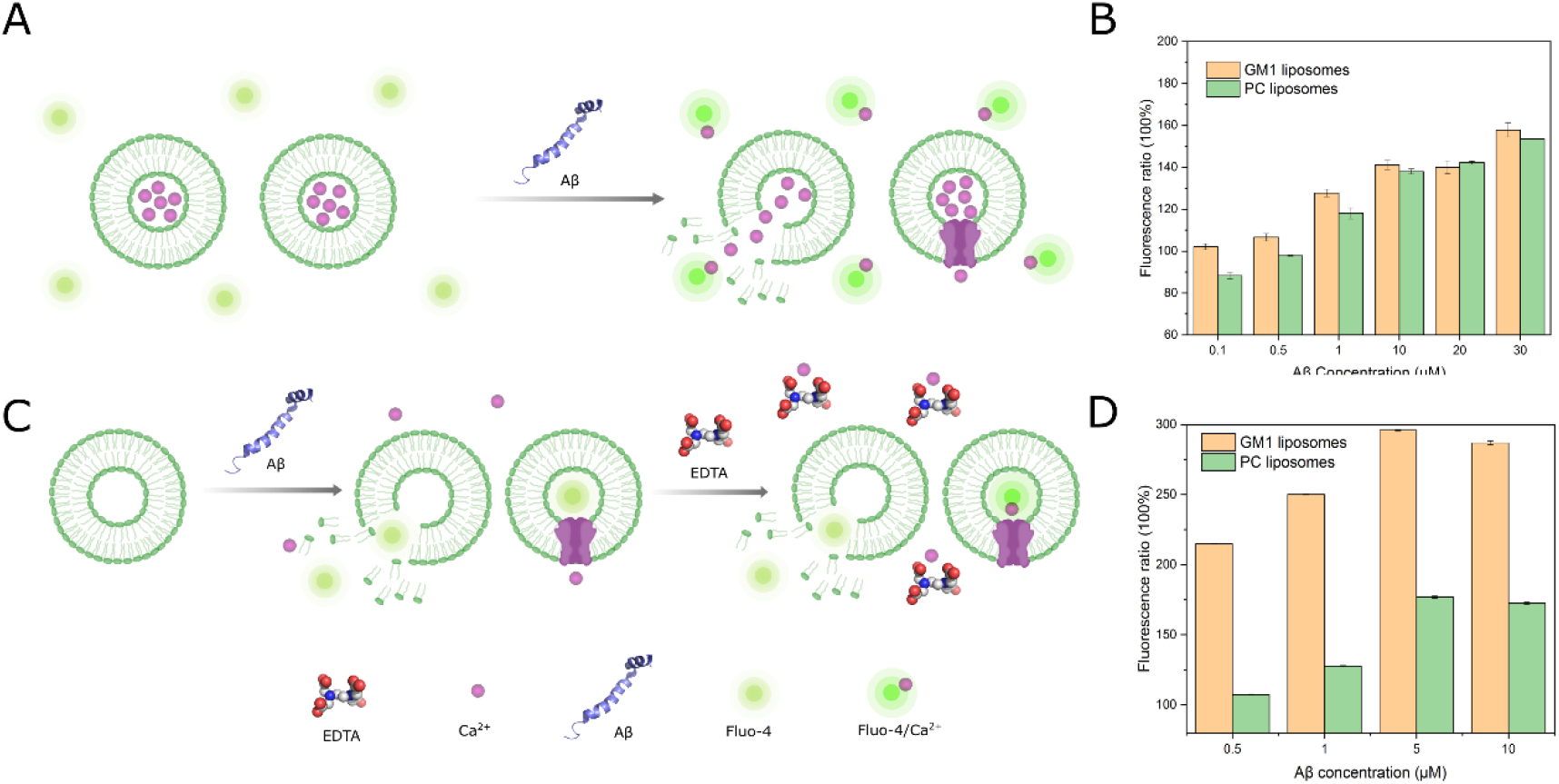
(A) Schematic of the experimental design to prove the formation of Aβ ion channels with Ca^2+^-encapsulated GM1 liposomes. (B) The ratio of the fluorescence intensity of Ca^2+^-encapsulated liposomes incubated with Aβ to that without Aβ. (C) Schematic of the experimental design to prove the formation of Aβ ion channels with dye-encapsulated GM1 liposomes. (D) The ratio of the fluorescence intensity of dye-encapsulated liposomes incubated with Aβ to that without Aβ.

We observe greater Ca^2+^ efflux in liposomes incubated with Aβ as compared to liposomes incubated without Aβ, suggesting that Ca^2+^ exits liposomes through two potential mechanisms: (i) Aβ forms channels on liposome membranes through which calcium can pass, or (ii) Aβ disrupts the integrity of the liposome membranes. To validate the formation of Aβ ion channels, we encapsulate Fluo-4 inside liposomes (Figure 4C&D) and incubate the liposomes with Aβ for 3 days. Next, we add extracellular Ca^2+^, incubate for another 3 h, and add an excess of ethylenediaminetetraacetic acid (EDTA) to chelate the Ca^2+^ outside the liposomes. EDTA has a higher binding affinity to Ca^2+^ than Fluo-4 (Park and Palmer, 2015). If Aβ disrupts the liposomes, the encapsulated Fluo-4 dye will diffuse to the solution outside of the liposomes, where it will be inert because all extracellular Ca^2+^ have been chelated by EDTA. Conversely, if ion channels are formed in the liposome membranes, Ca^2+^ will enter the liposomes through the channels to interact with Fluo-4 and emit fluorescent signals. Importantly, EDTA is too large to enter the liposomes through the ion channels. Indeed, we observe calcium influx, suggesting that Aβ forms ion channels capable of passing Ca^2+^ ions. Hence, we find that Aβ may form ion channels either spontaneously at high concentrations (~10 μM) or with GM1 as a catalyst at low concentration (~100 nM).

### Structural characterization of Aβ oligomers formed on membranes

To determine the size of Aβ oligomer species that can form channels, we utilize photo induced cross-linking of unmodified proteins (Fancy and Kodadek, 1999) (PICUP, Supplementary Figure 3) to cross-link Aβ oligomers and then remove Aβ oligomers from the solution using 1000 kDa MWCO centrifugation tube, so that only Aβ oligomers on the membrane are retained. We determine the molecular weight of the membrane-bound oligomers by mass spectrometry (MS) (Figure 5A&B). During PICUP, the formation of a cross-linking bond reduces the molecular weight by 2.016 Da, the mass of two hydrogens. We observe four possible molecular weights 18028.4, 72196, 85711.3, and 94711.9 Da, corresponding to 4, 16, 19, and 21 Aβ monomers with 14, 14, 27, and 41 cross-linkers, respectively. To determine the most probable number of monomers in Aβ oligomers that can form channels, we divide the oligomer samples into 4 parts according to the molecular weight: <25 kDa, 25-45 kDa, 45-100 kDa, and >100 kDa. We then incubate these four different Aβ samples with Ca^2+^-encapsulated liposomes and measure the fluorescence intensity, as described previously. The 45-100 kDa sample has the highest fluorescence intensity, indicating that the most likely molecular weight range of Aβ oligomers that can form channels is 45-100 kDa (Figure 5C), which may consist of 16, 19, or 21 Aβ monomers.

**Figure 5.**
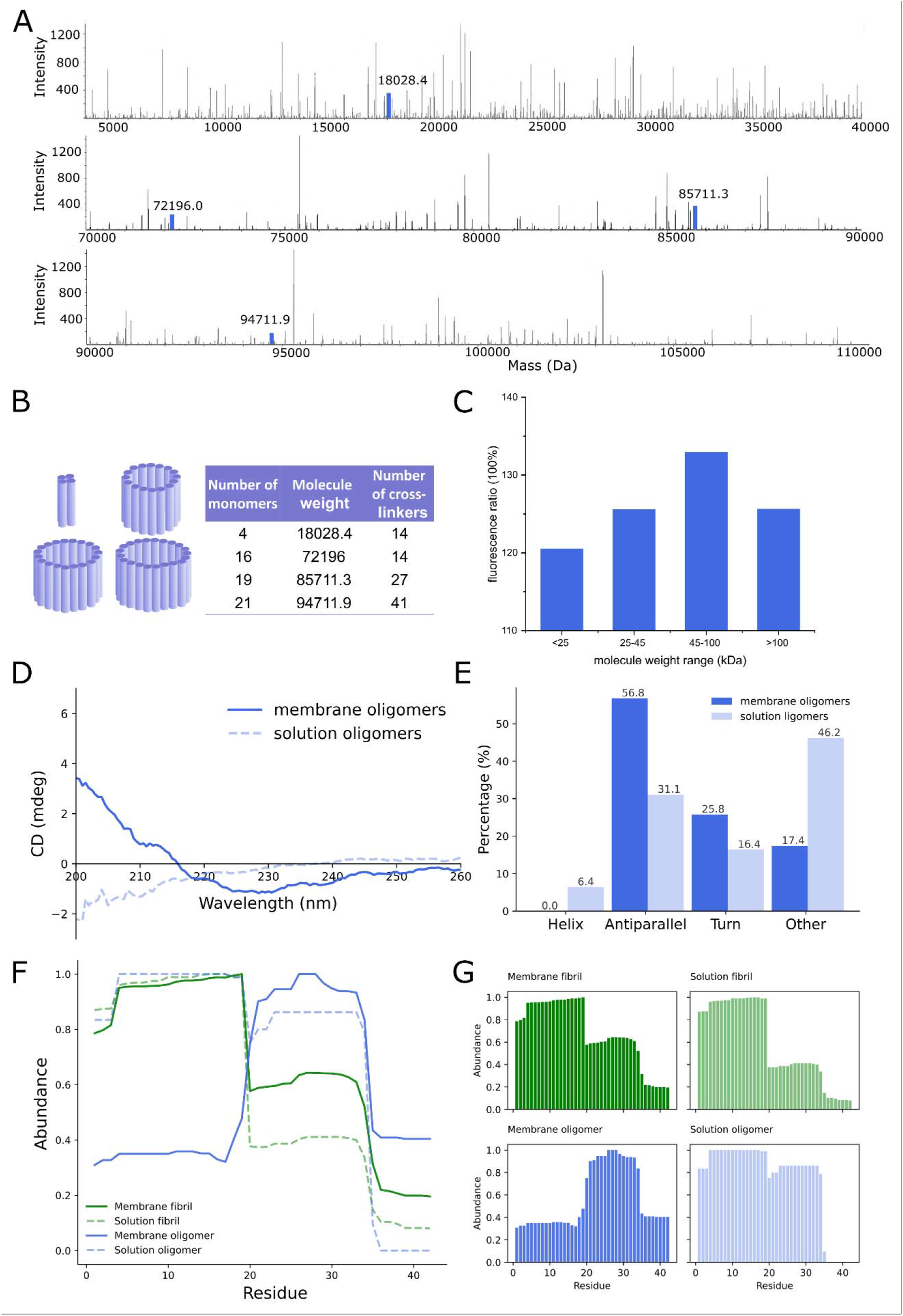
Structure characterization of membrane Aβ oligomers. (A) After purification, the molecule weights of Aβ oligomers are analyzed by mass spectrometer. (B) Based on the results of mass spectrometry, there are four possible cross-linked Aβ oligomer species that may form ion channels. (C) We divide the membrane Aβ oligomers into four samples according to the molecular weights. Then, we incubate the four samples with the Ca^2+^-encapsulated liposome and determine the fluorescence intensity. (D), (E) The secondary structure of the Aβ oligomers formed in the presence and absence of GM1 liposomes. (F), (G) The residue abundance of Aβ oligomers and fibrils formed in the presence and absence of GM1 liposomes.

We further determine the secondary structure of Aβ oligomers by circular dichroism (CD) and use BeStSel(Micsonai et al., 2018) to analyze the CD data in favor with its better estimation of β-sheet content. We prepare two oligomer samples, one obtained from the liposome membrane (membrane Aβ oligomers) and the other from the buffer solution (solution Aβ oligomers) (Figure 5D&E). Solution Aβ oligomers have 6.4% helices, 31.1% antiparallel beta-sheets, 16.4% turns, and 46.2% other structures, while membrane Aβ oligomers have a significantly higher percentage (56.8%) of antiparallel beta-sheet structures. Consistent with previous reports (Devarajan and Sharmila, 2014; Matsuzaki, 2014; Okada et al., 2007), our results show that GM1 membrane can promote a conformational change of oligomeric Aβ to form β-sheet-rich structures.

We compare the structural features of Aβ oligomers and fibrils by digesting all purified Aβ oligomers and fibrils with pepsin, and measuring the abundance of fragments by mass spectrometry (Proctor et al.) (Figure 5F&G). Theoretically, higher abundant peptides are likely to appear on the surface of the protein because surface residues have a higher chance to be digested, while lower abundant peptides are buried inside the structure (Proctor et al.). In both membrane Aβ fibrils and solution Aβ fibrils, the abundances of Aβ_4–19_ peptides are high, suggesting that they are on the surface; the abundances of Aβ_36–42_ peptides are low, suggesting that they are internal. Regardless of the presence or absence of GM1 liposomes, the abundances of peptides of the two fibrils are similar, indicating that although GM1 liposomes promote the formation of fibrils, they do not affect the structure of fibrils. However, the peptide abundances of the solution Aβ oligomers and membrane Aβ oligomers are drastically different. In membrane Aβ oligomers, Aβ_20–34_ peptides feature high abundances, while Aβ_1–17_ peptides feature low abundances. In solution Aβ oligomers, Aβ_4–19_ peptides have the highest abundances, and Aβ_35–42_ peptides have the lower abundances. Thus, the structures of membrane Aβ oligomers, promoted by GM1 liposomes, are distinct from the structures of solution Aβ oligomers. Finally, based on the abundances of peptides, the structures of Aβ oligomers and fibrils are also drastically different. Overall, the presence of GM1 liposomes changes Aβ oligomer structures but does not change Aβ fibril structures.

### Toxicity of Aβ oligomers *in vitro*

We interrogate the toxicity of these GM1-bound Aβ oligomers by cellular studies (Figure 6A). We incubate Aβ with GM1 liposomes, crosslink them, and remove solution Aβ oligomers. We then add the retained membrane Aβ oligomers to differentiated PC12 cells. We incubate Aβ with the cells for 3 days and then measure the toxicity by calcein AM and ethidium homodimer-1 live/dead viability assay. Membrane Aβ oligomers at the concentration of 5 μM exhibit toxicity (increased dead/live cells ratio) in PC12 cells (Figure 6B&C). While at the same concentration, solution Aβ oligomers do not exhibit toxicity (Supplementary Figure 4). Addition of membrane Aβ oligomers induces increased levels of cleaved caspase-3 compared to the live control (Supplementary Figure 5). The toxicity of membrane Aβ oligomers is consistent with our fluorescence results (Figure 4): the increase of Aβ oligomer concentration leads to the increase of Ca^2+^ efflux caused by membrane disruption, suggesting that the toxicity of membrane Aβ oligomers may be related to the dysregulation of calcium homeostasis.

**Figure 6.**
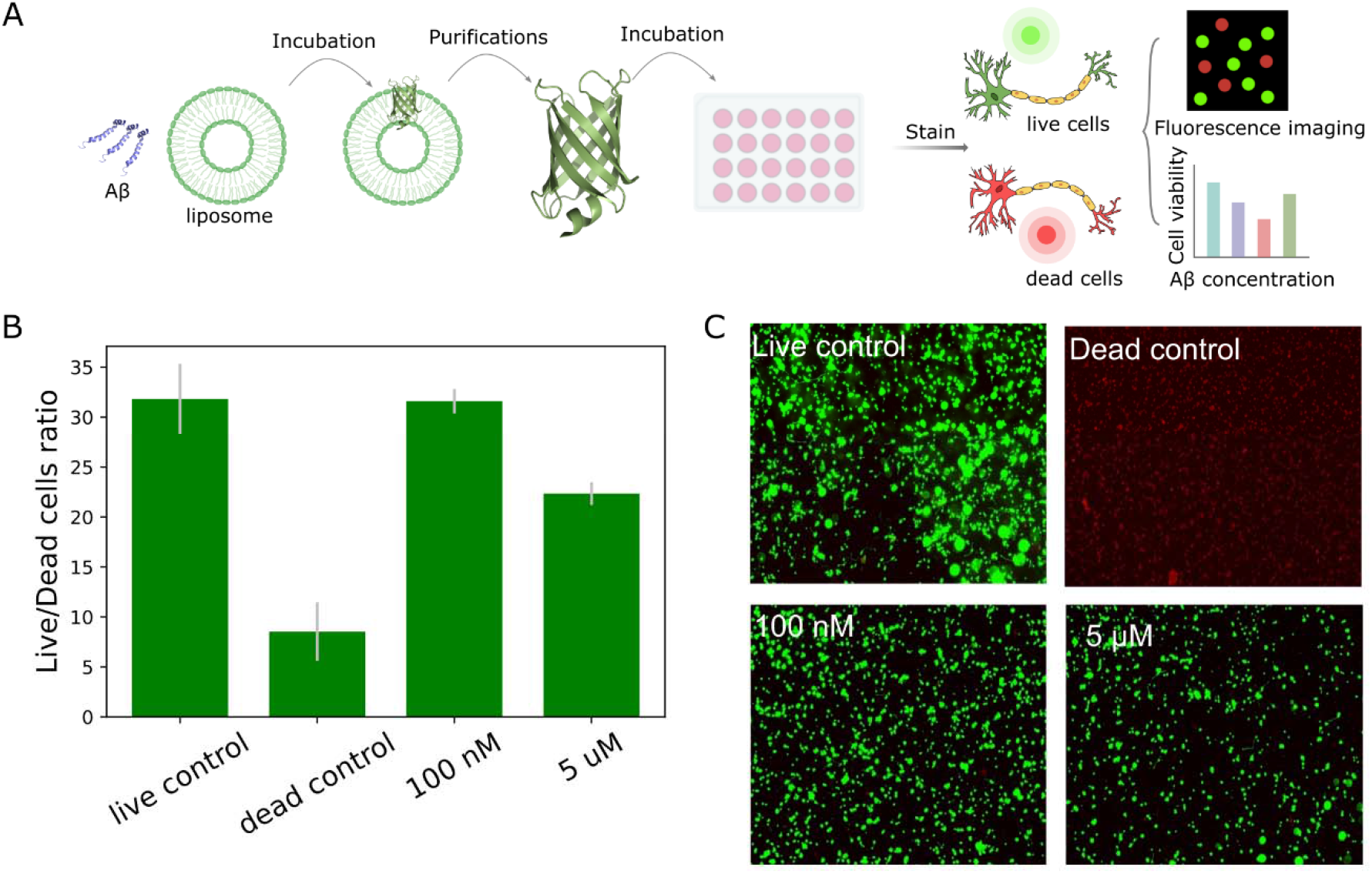
Aβ oligomers formed on liposome membranes are cytotoxic. (A) Schematic of cell viability experiment. (B) PC-12 cells incubated with and without Aβ oligomers. Cell viability is assessed by Calcein AM/ethidium homodimer-1 and calculated by the ratio of dead cells to live cells. (C) PC-12 cells incubated with Aβ oligomers of different concentration. Live cells are stained by Calcein AM in green and dead cells are stained by ethidium homodimer-1 in red.

Numerous GM1 gangliosides are present in the membrane of the differentiated PC 12 cell line (Nishio et al., 2004). To interrogate if membrane Aβ oligomers incur toxicity for cells with low GM1 content, we perform a viability assay using post-natal day 1 (P1) CD1 mouse cortical neurons, which have less GM1 since the amount of GM1 in the mouse brain has been previously reported to be minimal at birth, increasing with age (Yamamoto et al., 2008). Upon repeating the previously-described Fluo-4 assay in these primary neurons, we find that membrane Aβ oligomers do not result in calcium efflux (Supplementary Figure 6). The lack of effect on primary neurons, which are notoriously less robust than cell lines, suggests that the lack of GM1 in the cell membranes of the neonate-derived primary cells protects them from being compromised by Aβ oligomers, further supporting that GM1 is critically necessary for the toxicity of Aβ oligomers.

## DISCUSSION

The concentration of Aβ is important to the formation of oligomers (Hu et al., 2009; Thal et al., 2015). Most *in vitro* aggregation studies and Aβ ion channel studies have been conducted with Aβ peptide concentrations exceeding physiological concentrations by more than 1000-fold. Such a stark disparity between *in vitro* and physiological Aβ concentrations required for aggregation suggests the presence of an aggregation catalyst in living cells (Luo et al., 2013; Mantyh et al., 1993). Hence, the mechanism of oligomer formation at physiological concentrations remains one of the biggest mysteries of AD. Previous studies have demonstrated the destructive effects of Aβ on membranes with Aβ concentrations much higher than the physiological concentration (Chi et al., 2007). Here we demonstrate that phospholipid bilayer membranes can be disrupted by Aβ at nanomolar concentrations with GM1 serving as the catalyst.

GM1 is distributed as clusters on neuronal membranes in the mouse brain (Matsuzaki, 2014), and the binding of Aβ to GM1 clusters may increase the local concentration of Aβ and cause a conformational change that promotes aggregation (Fernández-Pérez et al., 2017; Ikeda et al., 2011; Yagi-Utsumi et al., 2010). The presence of GM1 clusters aiding the maintenance of Aβ oligomers may be due to the high binding affinity between GM1 clusters and Aβ. Previous reports and our results both suggest that GM1 can bind to Aβ (Kakio et al., 2002) and promote the formation of Aβ fibrils (Yanagisawa et al., 1995), and we further find that some Aβ oligomer species are stable and not inclined to further aggregate to fibrils in the presence of GM1 (Supplementary Figure 5). Our liposome encapsulation experiments demonstrate that these stable oligomers may form channel-like structures, which provides new evidences for the hypothesis that Aβ forms channel-like oligomeric structures that disrupt cellular calcium homeostasis (Jang et al., 2010). The formation of channel-like structures may be contributed by the conformational change of Aβ to β-sheet-rich states, which has been demonstrated in both previous studies (Chen et al., 2017; Matsuzaki, 2014) and our results. Thus, the mechanism of the disrupted calcium homeostasis may be that GM1 catalyzes the formation of Aβ ion channels thereby decreasing the calcium gradient (Ledeen and Wu, 2015). Furthermore, although some studies (Arispe et al., 1993b, 1993a; Jang et al., 2010) have utilized channel conductance measurements on planar lipid bilamellar membranes to suggest that Aβ forms ion channels, the size and structure information of the channel oligomers were unclear. In our work, we utilize mass spectrometry and CD to measure the possible size and secondary structure of Aβ oligomers, and we also find that the binding of Aβ to the GM1 membrane deforms the GM1 membrane.

Overall, we integrate computational, biochemical, and cellular studies to uncover the molecular etiology of Aβ aggregation modulated by GM1 in AD pathology and identify processes responsible for neuronal toxicity. Such a comprehensive approach is necessary to decouple multiple processes accompanying AD pathophysiology.

## METHODS

### Preparation of liposomes

We dissolve GM1 (Sigma-Aldrich), L-α-Phosphatidylcholine (PC, Sigma-Aldrich), sphingomyelin (VWR International), and cholesterol (Fisher Scientific) in a chloroform/methanol (1:1, v/v) mixture at a total concentration of 1 mM. To mimic the GM1 cluster on the cell membrane, the molar ratio of GM1, sphingomyelin, and cholesterol is 1:2:2 for GM1 liposomes. For PC liposomes, the molar ratio of PC, sphingomyelin, and cholesterol is 1:2:2. To prepare liposomes, we first dry the organic solvents under a gentle stream of nitrogen for 2 h and then under vacuum overnight. Then, we rehydrate the resulting lipid film with HEPES buffer (10mM HEPES and 150 mM NaCl, pH 7.4), and incubate 1 h at 45 °C water bath, vortexing every 15 min. In order to prepare Ca^2+^-encapsulated liposomes, we replace the rehydrated lipid HEPES buffer with 10 mM CaCl_2_ (Fisher Scientific), 10mM HEPES, and 140 mM NaCl, pH 7.4 (Colletier et al., 2002; Sanghera et al., 2011). To prepare dye-encapsulated liposomes, we replace the rehydrated lipid buffer with 0.03 mM Fluo-4 (Life Technologies), HEPES buffer. The dye-encapsulated liposomes are protected from light at all times. We sonicate the rehydrated suspension for 15 min, and then freeze-thaw for 5 cycles in the liquid nitrogen and 60°C water bath. To obtain uniform liposomes, we extrude the resulting suspension 16 times with an Avanti extruder. To encapsulate calcium and dye, we use 800 nm pore size polycarbonate filters to filter the liposomes. For the Ca^2+^-encapsulated and dye-encapsulated liposomes, we use 10000 Da molecular weight cut off (MWCO) dialysis cassette with HEPES buffer changed 3 times a day to remove the Ca^2+^ and dye in the buffer outside the liposomes.

### Fluorescence intensity determination

We dissolve Aβ42 (Fisher Scientific) in DMSO at a concentration of 1mM and sonicate for 10 min. For the Ca^2+^-encapsulated liposomes, we add different concentrations (100 nM, 500 nM, 1μM, 10 μM, 20 μM, 30 μM) of Aβ42 and incubate with liposomes for 3 days at room temperature. We add 50 μM Fluo-4 into the incubated Aβ42 and liposomes solution, and determine the fluorescence intensity using a SpectraMax i3 plate reader. We treat control sample in the same way, except without added Aβ42. We analyze the fluorescence intensity ratio of samples incubated with different concentrations of Aβ42 and control.

We protect the dye-encapsulated liposomes from light throughout the experiment to prevent photo-bleaching. We add different concentrations of Aβ42 (500 nM, 1 μM, 5 μM, 10 μM) into the liposome solutions and incubate for 3 days at room temperature. We add 14 mM CaCl_2_ into the incubated Aβ42 and liposomes and incubate for 1 h to ensure that calcium ions have enough time to enter the liposomes through the ion channels. Then, we add 16.5 mM EDTA (excess) to bind all the Ca^2+^ in the solution. In this way, only the Ca^2+^ that enters the liposomes can bind with Fluo-4. Similarly, we treat control sample in the same way, except without adding Aβ42. Finally, we determine the fluorescence intensity using a SpectraMax i3 plate reader. After incubation with GM1 liposomes, Aβ42 oligomers were divided into >100, 45-100, 25-45, and <25 kDa samples according to a trident pre-stained protein ladder (GTX50875) and then the same concentration and volume resulted sample was incubated with Ca^2+^-encapsulated GM1 liposomes overnight. After that, the fluorescence intensity was determined, as previously mentioned.

### Förster Resonance Energy Transfer (FRET)

We incubate 10 μM wild type or mutant Aβ42 with 200 μL GM1 liposomes for 3 days. After that, we add 10 μg/mL Alexa Fluor® 488 anti-β-Amyloid, 1-16 Antibody (6E10, Biolegend) to the incubated liposome solution to label Aβ42 as the donor, and 10 μg/mL Cholera Toxin Subunit B (Recombinant) Alexa Fluor™ 555 Conjugate (CTSB, Fisher Scientific) to label GM1 as the acceptor. We incubate Alexa Fluor® 488 6E10 and Alexa Fluor™ 555 CTSB with liposome solution for 1 h at room temperature. To remove the Alexa Fluor® 488 6E10 and Alexa Fluor™ 555 CTSB that did not bind to Aβ and GM1 liposomes, we use 1000 kDa MWCO centrifugation tube at 1000 rpm and dilute samples 1:40 in HEPES buffer. and centrifuge to the final volume of 50 μL and repeat this process. We use SpectraMax i3 to analyze the labeled sample. The excitation wavelength is 492 nm with 2 nm slit width. We collect the emission spectra from 518 to 750 nm. For the donor control, we treat sample in the same way except we do not add the acceptor (Alexa Fluor™ 555 CTSB) to label GM1. For the acceptor control, we treat sample in the same way except without Alexa Fluor® 488 6E10 to label Aβ.

### Fluorescence characteristics of Aβ fibrils using Thioflavin T (ThT)

We determine the effect of GM1 liposomes on Aβ aggregation using ThT (VWR International). We incubate 80 μM wild type or mutant Aβ monomers in the presence or absence of GM1 liposomes containing 10 μM ThT at 37 ^o^C shaking in black 96-well plates with seal. We treat HEPES buffer containing 10 μM ThT in the presence and absence of GM1 as control. We collect the fluorescence intensities of all samples every 30 seconds for 3 days using a SpectraMax i3 plate reader with excitation wavelength of 445 nm and emission wavelength of 490 nm.

### Cross-linking

To obtain stable Aβ oligomers, particularly the Aβ oligomers formed on membranes, we utilize the photo-induced cross-linking of unmodified proteins (PICUP) method (Bitan and Teplow, 2004) to cross-link. We add ammonium persulfate (APS 1 mM, 10μL, Dot Scientific) and Tris (2,2-bipyridyl)dichlororuthenium(II) hexahydrate (Ru(II)Bpy_3_^2+^ 0.05 mM, 10 μL, Sigma-Aldrich) in 10 mM HEPES (pH 7.4) to the incubated Aβ oligomer solution (180 μL). We irradiate the resulting solution with a 500 W lamp for 5 seconds and quench the cross-linking process with 47.6 mM dithiothreitol (DTT 1M, 10 μL, Dot Scientific).

### Analysis of the size of Aβ oligomers by SDS-PAGE and silver staining

We incubate Aβ in the presence or absence of GM1 liposomes and analyze samples at 0.5 h, 4 h, 24 h, and 72 h. We cross-link all samples using the PICUP method. For the sample in the presence of GM1 liposomes, we add 40 μM N-octylglucoside (Dot Scientific) into the samples and then incubate at room temperature for 1 h to dissolve the lipids of the liposomes. We add leammli SDS sample buffer dye (Fisher Scientific) to the resulting solution and heat at 95°C for 10 min. We use 10% SDS-PADE and silver staining (Fisher Scientific) to analyze the formed Aβ oligomers.

### Purification of Aβ42 oligomers on the liposomes

We incubate Aβ42 monomers with GM1 liposomes at room temperature for 3 days and cross-link Aβ before further purification. We use 1000 kDa MWCO centrifugation tubes, spun at 1300 rpm, to remove Aβ oligomers that did not insert into liposomes. We treat sample with N-octylglucoside, leammli SDS sample buffer dye, heater, SDS-PAGE, and silver staining as described above. After gel electrophoresis, we cut out the gel portion according to the reference results of silver staining. We crush the cut gel portion containing the Aβ oligomers of interested and dissolve them in HEPES buffer overnight in a shaker at 4°C. We centrifuge the resulting samples at 10000 g for 10 min and save the supernatants for further concentration by centrifugation. To measure the size of the Aβ channels, we cut the gel according to various molecular weight ranges (>100, 45-100, 25-45, <25 kDa) and purify Aβ oligomers in the gel as described above. We determine the concentration of purified Aβ sample using Nanodrop and a bicinchoninic acid (BCA) protein assay kit.

### Secondary structure

We incubate Aβ42 monomers (10 μM) for 3 days at room temperature in the presence or absence of GM1 liposomes, and then cross-link samples using PICUP. For the sample of Aβ42 incubated in the presence of GM1 liposomes, we remove Aβ oligomers in the solution and add N-octylglucoside to dissolve the lipids as described previously as mentioned previously. We remove the fibrils in the resulting solution using 1000 kDa MWCO centrifugation tubes spun at 3900 rpm, and wash with HEPES buffer. We remove the monomers, dimers, and lipids using 10 kDa MWCO centrifugation tubes, spun at 3900 rpm, and wash with HEPES buffer with 40 μM N-octylglucoside and then wash again with HEPES buffer. For the control sample of Aβ42 monomers (10 μM) incubated in HEPES without liposomes, we remove the fibrils, monomers, and dimers in the same way except adding N-octylglucoside to dissolve the lipids and washing with 40 μM N-octylglucoside HEPES buffer. We determine the concentration of Aβ oligomers using BCA protein assay kit. We determine the secondary structure characteristics of Aβ42 channels and oligomers using a Jasco J-1500 circular dichroism spectrophotometer. We place samples in 0.1-mm path-length (200 uL) cuvettes, and record the spectra range at 185-240 nm with 0.5 nm data pitch and scanning speed of 50 nm/min. We scan each sample three times and calculate average. We use the software BESTSEL(Micsonai et al., 2018) to analyze experimental data and get the fractional content of the helix, sheet, turn, and other structures element.

### PC-12 cell culture

We obtain PC-12 cells from the American Type Culture Collection (ATCC). For the PC-12 cells, we use DMEM (VWR International) supplemented with 5% heat inactivated horse serum (Life Technologies), 5% fetal bovine serum (Fisher Scientific), and 1% Penicillin/Streptomycin (Fisher Scientific). We culture cells at 37°C with 5% CO_2_. For the neuronal differentiation, we seed PC-12 cells in 96-well plates at 2000 cells/well. Prior to seeding, we treat the 96-well plate by UVO-cleaner (Jelight Company, model 18) for 30 min, coat with 10X diluted collagen and incubate at 4°C for 2 h before used. For the differentiated PC-12 cells, we use DMEM supplemented with 1% heat inactivated horse serum, 1% Penicillin/Streptomycin, and 100 ng/mL nerve growth factor (Sigma Aldrich) and culture for 1 week at 37°C with 5% CO_2_. Before further analysis, we confirm differentiation by visual observation using a phase contrast microscope.

### Cell viability assay

We determine cell viability using calcein AM and ethidium homodimer-1 dye. After differentiation, we add purified Aβ oligomers samples to the differentiation cells and incubate for 3 days. For the dead cell controls, we add 70% methanol to cells 2 h before plate read. We remove the differentiation medium and wash the cells twice with PBS. For each well, we add 4 μM ethidium homodimer-1 and 2 μM calcein AM in PBS to label live and dead cells. Cells then incubate for 30 min at room temperature. We determine the fluorescence intensity by SpectraMax i3. For calcein AM, the excitation wavelength is 495 nm and the emission wavelength is 525 nm. For ethidium homodimer-1, the excitation wavelength is 495 nm and the emission wavelength is 645 nm. We take the fluorescent images on a Keyence Bz-x800 fluorescent microscope.

### Western blot

We incubate the differentiated PC-12 cells in the presence and absence of purified Aβ oligomer samples for 3 days and then lyse with ice-cold RIPA buffer (PBS, 1% Nonidet P-40, 0.5% sodium deoxycholate, 0.1% SDS). We then collect the lysed solution for Western blot. After homogenization, we centrifuge cell samples at 12000 rpm for 20 min and collect the supernatants. We add 2X SDS sample buffer into the supernatant solutions and heat samples at 95 °C for 10 min. We separate the samples on a 12% polyacrylamide gel then transfer to a PVDF membrane with transfer buffer (25mM Tris, 192mM Glycine, 0.1% SDS,20% Methanol, pH 8.3) at 80 V for 90 min at 4°C. We block the blots with 5% BSA in TBST (10 mM Tris-HCl, pH 7.6, 150 mM NaCl, 0.1% Tween-20) for 1 h at room temperature, then incubate with anti-Caspase 3 (Sigma AB3623) and anti-Vinculin (Sigma SAB4200729), diluted 1:100 and 1:1000, respectively, in blocking buffer (caspase-3 and vinculin) overnight at 4°C. After washing 3 times with TBST, we incubate the membrane in anti-Mouse IgG conjugated with horseradish peroxidase for 3 hr at room temperature. Before film imaging, we wash the blot 3 times with TBST and soak for 5 min in chemiluminescent substrate (Thermo Scientific catalog # 34080).

### Mass Spectrometry (MS)

After cross-linking and purification, we measure the molecular weight of the Aβ oligomers with MS using the Sciex 5600 triple tof (AB SCIEX). Then, we digest purified Aβ oligomers and fibrils with pepsin. We add 1 N HCl to the Aβ samples to a final concentration of 0.04 N and suspend pepsin in 100 mM acetate buffer (pH 3.5). We add the pepsin to Aβ solution as the ratio of 1:20 (enzyme:protein W:W) and incubate them for 1 h at 37°C. We add 1 M Tris buffer (pH 8) to a final concentration of 150 mM to stop the reaction. And then we determine the resulting peptides with Sciex 5600 triple tof (AB SCIEX) and analyze them using the ProteinPilot software.

### Cryo Electron Tomography

To increase the ratio of Aβ:lipid to 1:10, we dilute GM1 liposomes 10 times and incubate the resulting GM1 liposomes with or without 10 μM Aβ for 3 days. Then, we concentrate the samples using a 100 kDa MWCO centrifuge tube, centrifuging at 1300 rpm to reduce sample volume from 1 mL to 50 μL to increase the concentration of liposomes. We mix the samples with 10 nm gold fiducials before pipetting 4 μl onto a freshly glow-discharged Quantifoil R2/2 holey carbon grid. We hand grids blotted from behind and plunge grids into liquid ethane using a Mark IV Vitrobot (Thermo Fisher Scientific) for vitrification. We then transfer samples under liquid nitrogen into a Titan Krios G3i cryo TEM (Thermo Fisher Scientific) operating at 300 kV for the acquisition of tilt series. We target holes containing liposomes contained within vitreous ice data collection and collect tilt series in 2° increments from −60° to + 60° at −6.0 μm defocus. Tilt series images are acquired with a Bioquantum energy filter (Gatan) operating at the zero-loss peak using a K3 direct electron detector (Gatan) in single electron counting mode. We use a nominal magnification of ×33,000 which correspond to a pixel size of 3.3 A◻/pixel. We set exposure time to achieve a total electron dose of ~120 e^−^/A◻^2^ for a complete tilt series. We collect six to ten tilt series from each grid. We reconstruct tomograms using the IMOD(Kremer et al., 1996) software suite.

### Computational modeling

We first model the GM1 membrane structure through CharmmGUI (Jo et al., 2008) with the ratio of SM, Cholesterol, GM1 as 4:4:2. We then model the intact structure of Aβ using SwissModel (Schwede et al., 2003) with PDB ID 1IYT as the template. The missing residues in the N-terminal region are completed by SwissModel. Next, we perform molecular dynamics simulation of the GM1 membrane-Aβ system using GROMACS (Van Der Spoel et al., 2005). The Charmm36m (Huang et al., 2017) force field for the protein and lipid parameters are used. Hydrogens for heavy atoms were added by the pdb2gmx module in the GROMACS simulation package. The system is subsequently energy-minimized for 2000 steps using the conjugate gradient algorithm, and another 2000 steps using the steepest descent algorithm. The structure is solvated using explicit water in a cubic periodic box with water molecules extending 10 Å outside the protein on all sides. Water molecules are described using a simple point charge water model. The system is solvated using TIP3 water molecules, then energy-minimized again and heated up gradually to reach a temperature of 310 K using a V-rescale thermostat with a coupling constant of 0.1 ps. The solvent density is adjusted under isobaric and isothermal conditions at 1 bar and 310 K. A Parrinello-Rahman barostat with isotropic pressure coupling and a coupling constant of 0.1 ps was used to set the pressure at 1 bar. The system is equilibrated for 10 ns in the NPT ensemble with a simulation time step of 2 fs. Finally, the production run is carried out for 100 ns for the primary system. The long-range electrostatic interactions are treated using particle-mesh Ewald sum with a cut-off of 1.0 nm. The van der Waals interactions are terminated beyond the cut-off value of 1.0 nm. The LINCS algorithm is used to constrain all bonds involving hydrogen atoms. All simulations were performed using the GROMACS simulation program. The analyses of the trajectories are performed using GROMACS and MDAnalysis (Michaud-Agrawal et al., 2011)

## ACKNOWLEDGEMENTS

We acknowledge support from the National Institutes for Health 1R35 GM134864, the Huck Institutes of the Life Sciences, and the Passan Foundation. The project described was also supported by the National Center for Advancing Translational Sciences, National Institutes of Health, through Grant UL1 TR002014. The content is solely the responsibility of the authors and does not necessarily represent the official views of the NIH. We also acknowledge the help from Brianna L. Hnath, Martin Dokholyan and Sophia Dokholyan with the purification of membrane Aβ oligomers.

## Notes

### Competing Interest Statement

The authors have declared no competing interest.

