## Supplemental information for "GM1 mediates the formation and maintenance of cytotoxic Aβ oligomers"

#### Supplemental Methods

##### Primary neuron culture

We use postnatal day 0/1 (P0/1) CD1 pups for the generation of primary neuron cultures. CD1 breeder pairs (Charles River) are fed food and water *ad libitum* and maintained under 12 hr light/dark conditions under a protocol approved by the Penn State College of Medicine Institutional Animal Care and Use Committee (IACUC). All procedures are conducted according to NIH and approved IACUC guidelines. For the generation of primary neuron cultures, we sacrifice P0/1 pups by decapitation using surgical scissors. We place isolated brains in cold HEPES buffered Hanks' Balanced Salt solution (pH 7.8). We dissect out the cortical cap of each brain and remove the meninges. Isolated cortices are transferred to conical tubes of warm neuron plating medium: Neurobasal Plus (Gibco), 10% FBS (Gibco), 1x GlutaMAX (Gibco), 1x Penicillin-Streptomycin (10,000 U/mL, Gibco). We triturate ortices in plating medium with a cut p1000 pipette tip, followed by trituration with an intact tip. We measure the cell concentration using the Countess II automated cell counter (Invitrogen), and plate 5.47*105 cells/$cm^2$ on poly-D-lysine (Cultrex)-coated 96-well plates. We place the cells in the incubator at 37ºC, 5% $CO_2$ overnight to allow cells to attach. The next morning, we switch the medium to Neuronal Medium: Neurobasal Plus (Gibco), 1X B27 Plus supplement (Gibco), 1x GlutaMAX (Gibco), 1x Penicillin-Streptomycin (10,000 U/mL, Gibco). We change half the medium every 5-7 days and used confluent neuron cultures for calcium and viability assays after 12-14 days in culture.

##### Measurement of $Ca^{2+}$ influx in primary neurons

We solubilize human Aβ42 (Novex 75492034A) in trifluoroacetic acid to a concentration of 110 μM and vortex vigorously. We initiate Aβ42 aggregation by diluting the peptide solution to 11 μM in PBS, and incubate at 37 ºC for 24 hours. As a control, we dilute the same volume of TFA vehicle in PBS and incubate at 37 ºC for 24 hours. We then dilute each of the aggregated Aβ42 and TFA control in neuronal media to the indicated concentrations. We remove all media from the neurons, and replace with the Aβ42 media or TFA vehicle control media. After 72 hours, we wash cells twice with artificial cerebrospinal fluid (aCSF) (124mM NaCl, 4.4 mM KCl, 1.2mM $MgSO_4$:7$H_2O$, 1mM $NaH_2$PO4:$H_2O$, 2.175g NaHCO3, 2.5mM $CaCl_2$:2$H_2O$, and 1.825g glucose in 1L of dH2O, pH 7.4). We then add 100uL of 2.3uM Fluo-4 (diluted in aCSF) to each well and incubate at 37ºC to allow Fluo-4 to enter the cells. After 15 minutes, we aspirate the wells and wash twice with calcium-free aCSF. We add 100uL of aCSF to all wells, except a set of wells which receive 10 uM ionomycin (a membrane permeable calcium ionophore) as a positive control. We then determine fluorescence intensity using a SpectraMax i3 plate reader.

### Supplemental Figures

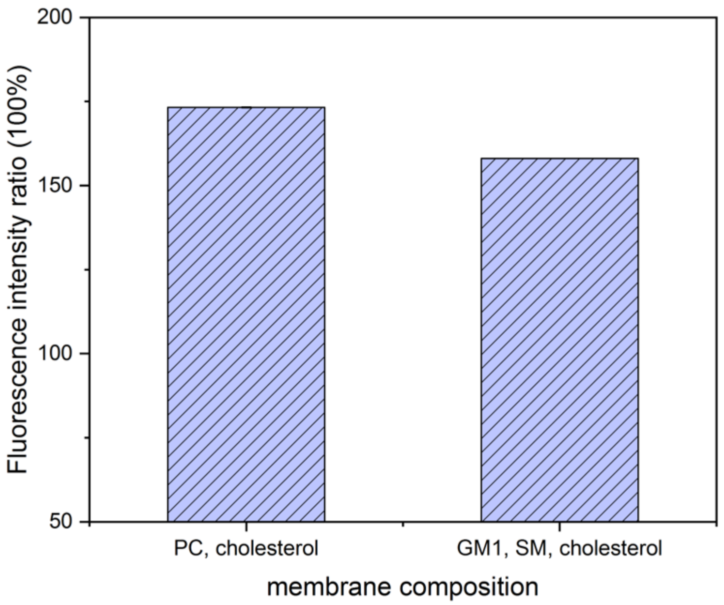


**Figure S1**. **Validation of the encapsulation of $Ca^{2+}$ in liposomes.** The y-axis is the ratio of the fluorescence intensity of $Ca^{2+}$-encapsulated liposomes treated by Triton X-100 to the fluorescence intensity of $Ca^{2+}$-encapsulated liposomes not treated by Triton X-100. Triton X-100 could dissolve lipid and destroy liposomes, thereby releasing $Ca^{2+}$ encapsulated in the liposomes. The ratios of both PC/SM/cholesterol and GM1/SM/cholesterol membranes are higher than 100%, suggesting that $Ca^{2+}$ are successfully encapsulated in liposomes.

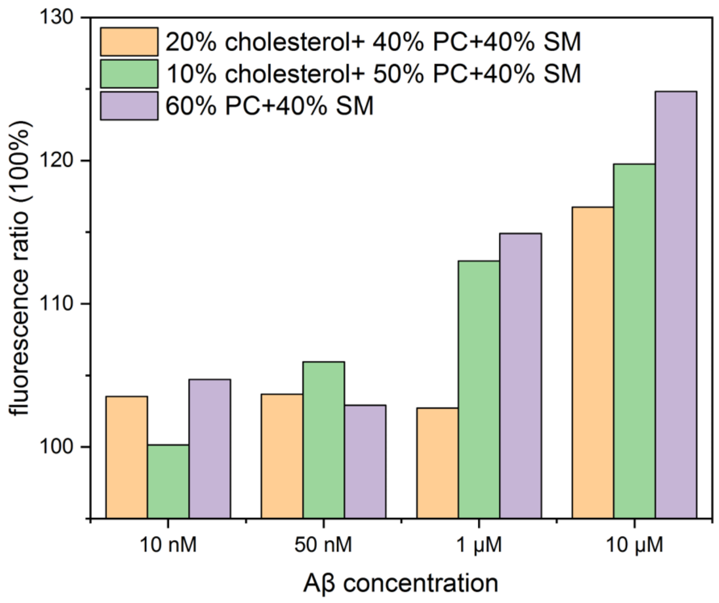

**Figure S2**. **The effect of different lipid compositions (PC, SM, and cholesterol) on Aβ destroying $Ca^{2+}$-encapsulated liposomes.** We prepare $Ca^{2+}$-encapsulated liposomes of different lipid compositions and Aβ of different concentration. We then incubate them and determine the ratio of fluorescence intensities of liposomes incubated with Aβ to the fluorescence intensities of liposomes incubated without Aβ. We find that the change of PC, SM, and cholesterol content in liposomes have no significant effect on the ratio of the fluorescence intensity.

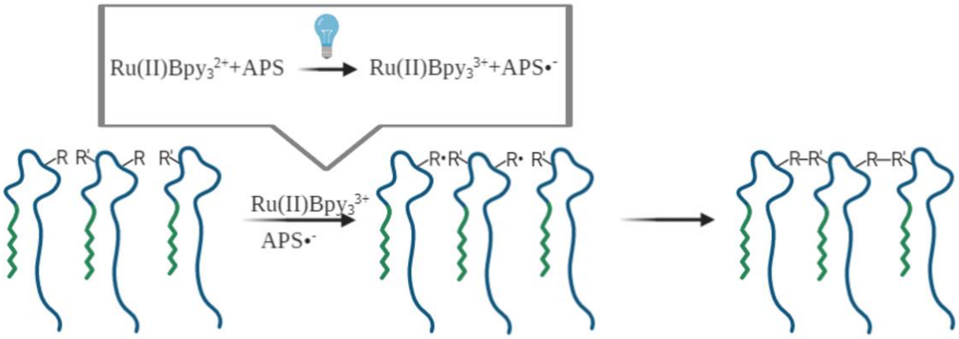


**Figure S3**. **The schema of PICUP reaction mechanism**. R is the reactive group in Aβ. R' is the adjacent reactive side-chains in Aβ.

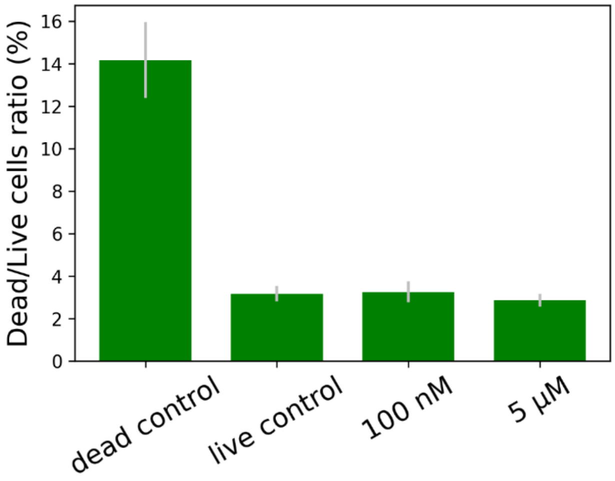


**Figure S4.** The toxicity of solution Aβ oligomers**.** PC-12 cells are incubated with and without solution Aβ oligomers. Cell viability is assessed by Calcein AM/ethidium homodimer-1 and calculated as the ratio of dead cells to live cells.

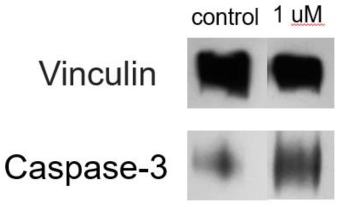


**Figure S5. The levels of the apoptotic marker cleaved caspase-3 and vinculin in PC-12 cells.** Vinculin is used as the control. PC-12 cells are incubated in the presence or absence of membrane Aβ oligomers. The level of Caspase-3 is measured by western blot.

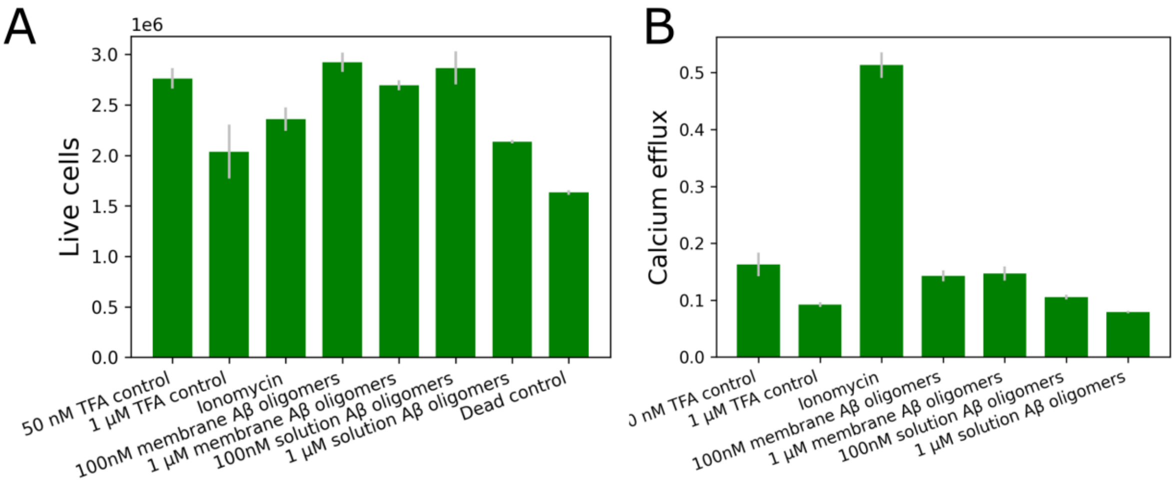


**Figure S6. Cell viability and calcium efflux in the primary neurons.** (A) Characterization of the cell viability of primary neuron cells. The treatment of TFA vehicle and methanol are served as the live control and dead control. (B) We calculate the ratio of the determined fluorescence intensity to the corresponding live cells as the final calcium efflux.
